# HyPhy: a skeletonization-based approach for fungal network analysis

**DOI:** 10.1101/2025.09.11.675604

**Authors:** Melanie Madrigal, Aaron J. Moseley, Jenna C. Moseley, Jordan A. Dowell

## Abstract

**Premise:** Traditional methods to quantify mycelial growth rely on destructive sampling to quantify biomass. However, these approaches limit continuous observation and require a large enough mass to measure. Recent work examines hyphal network traits by reconstructing the hyphal network from spatial coordinates via images, providing information about branching patterns and spatial growth over time.

**Methods and Results:** We developed HyPhy, a Python-based graphical user interface that skeletonizes images of hyphal networks and extracts biologically relevant structural parameters such as fractal dimension, a proxy for the complexity and branching structure of the hyphal network. Using a high-throughput pipeline method, we imaged three isolates of *Botrytis cinerea* grown under liquid culture for 72 hours, generating a dataset of 180 time series images.

**Conclusions:** HyPhy enables efficient, non-destructive, and scalable quantification of hyphal growth and complexity from time-resolved image datasets, providing a powerful and user-friendly tool for studying fungal network dynamics.

## INTRODUCTION

Fungi are eukaryotic organisms that play several roles in our environments, spanning decomposers that help break down organic matter to destructive pathogens. Fungi can exist as unicellular organisms like yeast or through multicellular networks. The fungal body is composed of hyphae which function as the major building block of fungi. Hyphae are characterized by their thin, tubular structures that grow via apical extensions and branching to maximize foraging and nutrient acquisition. As hyphae expand, they interconnect and form complex hyphal networks that are fundamental to development and environmental interactions. Hyphal growth is largely driven by the number of active tips; hyphal tips house the Spitzenkorper. The Spitzenkorper acts as a vesicle supply center and generates a polarity feedback loop that drives tip extension (Steinberg et al., 2017).

As heterotrophs, fungi rely on absorbing nutrients via their surroundings. Under low-nutrient conditions hyphae exhibit exploratory growth, characterized by the formation of sparse outward hyphae, with spaced branches, that search for nutrients. In contrast, under high nutrient availability, fungi exhibit invasive growth, characterized by compact multi-directional branching that allows for denser colonization. The plasticity of hyphal networks suggests that quantifying variation among these networks may provide insight into foraging strategies and how strategies may shift in response to environmental changes. In addition to variation in network density, some fungal pathogens form sclerotia primordia, which are solid-like, dense, and armored hyphal clusters that form in response to stressful environmental changes (Ordóñez-Valencia et al., 2015). Thus, variation in the size and pattern sclerotia primordia may indicate the degree of environmental stress. Therefore, analysis of fungal structural changes via image analysis may lead to novel biological insight combing foraging strategy from overall network parameters and severity of environmental stress via variation in sclerotia primordia.

Traditionally to quantify hyphal length density, the grid line method is used where a grid is placed over an image of hyphae, and the intersections of the hyphae are used to calculate hyphal length density as described by (Marsh, 1971; Newman, 1966). This, however, requires time, and the reliance on user dependance quantification introduces bias. To minimize user bias and labor, (Vidal-Diez De Ulzurrun et al., 2015) developed an automated image analysis software. Vidal-Diez de ulzurrun et al., collected images of agar droplets on petri dishes using a flatbed scanner, imaging every 30 minutes for 72 hours. The culmination of their work was graphical user interface (GUI) that extracts hyphal network parameters (Vidal-Diez De Ulzurrun et al., 2015) by binarizing images of samples, skeletonizing these images, and leveraging graph theory to quantify morphological traits such as fungal network growth, hyphal length, and hyphal tip width. However, these applications are developed for MATLAB which is not a free open-source software, limiting accessibility. Additionally, the experimental setup described Vidal-Diez de Ulzurrun et al., only allows for 6 samples to be imaged simultaneously, limiting the scalability of experiments that might require imaging hundreds of samples simultaneously. In this study, we used a high-throughput method where 3 isolates of *Botrytis cinerea* were grown under liquid culture in a 96-well microplate for 72 hours and imaged every four hours using a CytationGen5 plate reader (Agilent Technologies, Santa Clara, California, USA), generating a dataset of 180 time series images. These images were used to develop HyPhy a user-based software compatible-GUI that allows users to change settings to maximize image processing to deliver skeletonized images, which can be used for downstream analysis to calculate traits described by Vidal-Diez de Ulzurrun et al., such as the number of tips, edges, and nodes.

## METHODS AND RESULTS

HyPhy is an open-source graphical user interface program written in Python3 using PyQt. The source repository and user instructions can be found at https://github.com/AaronMoseley/HyPhy.git. The software can be easily installed on Windows, OS X, and Linux by cloning the repository. Before use, users need to set up a conda environment and install the program dependencies. Detailed instructions on the installation of HyPhy can be found on the readme page of the GitHub repository.

### Dataset

Three isolates of *B. cinerea* (KatieTomato, B05.10 and 1.04.25) that span in virulence (Caseys et al., 2021) and genetic diversity (Atwell et al., 2018) were cultured for two weeks on oatmeal agar plates. KatieTomato is a highly virulent isolate, characterized aggressive infection and the formation of large lesions on its host. B05.10 represents an intermediate-virulence isolates, while 1.04.25 is a slow growing isolate with reduced pathogenicity.

Spores were collected using sterile water and diluted to a final concentration of 10 spores/µL using an automated spore counting pipeline in R, which can be found at https://github.com/melaniemadrigal07/HyPhyAnalysis.git. Spores were kept on ice from collection to inoculation to prevent spore germination. Spores were then inoculated in to a 96-well microplate with 70 µL of liquid gamborg B5 media (Research Products International, Mount Prospect, Illinois, CA) (supplemented with 41 grams fructose, 26.65 grams glucose, 1.03 grams sucrose, pH= adjusted to 6.5), 10 µL of 1000 uM Linalool (Thermo Fisher Scientific, Waltham, Massachussets, USA), 10 µL of spore suspension (10 spores/µL), and 10 µL of OZ blue cell viability (OZBiosciences, Marseille, France), in triplicate. Linalool was included as part of a larger study of resistance among *Botrytis* isolates to linalool and a subset of that studies images were used to develop this method.

### Image Capture

Plates were incubated and imaged every four hours across 20 timepoints (n=180) using a Biotek Cytation Gen5 Plate Reader within a BioSpa session. Color brightfield images were captured at 4X magnification for each color channel (RGB) and exported as individual .TIF files. For each timepoint, the channel-specific images were stacked into three-channel composite images using a python script available in the HyPhyAnalysis repository (https://github.com/melaniemadrigal07/HyPhyAnalysis.git). To facilitate user familiarity with the workflow, the HyPhy repository includes a subset of images for users to practice with HyPhy.

### User Interface

Upon running the program, users are prompted to select an input directory containing grayscale experimental images in .tif/.png format, this ensures compatibility with HyPhy. For compatibility, files must follow the pattern {sample name}_timestep.{.png or .tif}. An output directory must also be provided for saving the processed images (**Fig. 1**). Users can generate skeletons for all images in the directory, a single sample timepoint course, or a specific image to skeletonize. After running HyPhy, images can be viewed with preset thresholds and modify the pipeline parameters such as the gaussian blur, and noise filtering to optimize skeleton generation of each sample, which are described in **Table 1** for sclerotia primordia and hyphal network parameters.

**Table 1.** Pipeline Parameters.

| Processing Step<sup>a</sup> | Description | Increasing | Decreasing |
| --- | --- | --- | --- |
| **Radial Thresholding<sup>b</sup>** | Creates threshold mask that circularly interpolates from center threshold at the center to the edge threshold at the edges. | Center Threshold: Central pixels appear white<br>Edge Threshold: Edge pixels appear black | Center Threshold: Central pixels will appear white<br>Edge Threshold: Edge pixels appear black |
| **Remove Small White Islands<sup>c</sup>** | Counts the number of pixels in each group of 1's.<br>Area with fewer pixels than this parameter turn black. | Removes more white areas; larger areas remain. | Preserves white areas; increases speckles in the image. |
| **Remove Noisy Islands<sup>d</sup>** | Counts the average number of black neighbors in each connected group of pixels. If average is greater than the threshold, area turns black. | More areas are considered noise; fewer white islands remain. | Fewer areas are noise; more white islands are present. |
| **Smooth Image<sup>e</sup>** | Reduces image noise and detail using a Gaussian Blur, helps enhance image structures | Smoothness increases; reduces fine detail. | Weaker smoothing; keeps more detailed structures but also increases noise present. |
| **Adjust Contrast<sup>f</sup>** | Scales pixel values away or toward 0.5.<br>Used to easily identify network lines | Scales pixels away from 0.5<br>Enhances network lines, preserves interconnect. | Scales pixels toward 0.5<br>Reduces contrast, network lines are less preserved. |
| **Edge Detection<sup>g</sup>** | Uses Canny Edge detection to outline the fungal network. | Detects more edges, but can increase noise. | Detects fewer edges. Faint structures may be missed. |

*Note:*<sup>a</sup>All processing steps are part of the HyPhy pipeline.<sup>b</sup>Parameters = centerThreshold, edgeThreshold<sup>c</sup>Parameters = minWhiteIslandSize<sup>d</sup>Parameters = noiseTolerance<sup>e</sup>Parameters = gaussianBlurSigma
<sup>f</sup>Parameters = contrastAdjustment
<sup>g</sup>Parameters = gaussianBlurSigma, maxThreshold.

**Figure 1.**
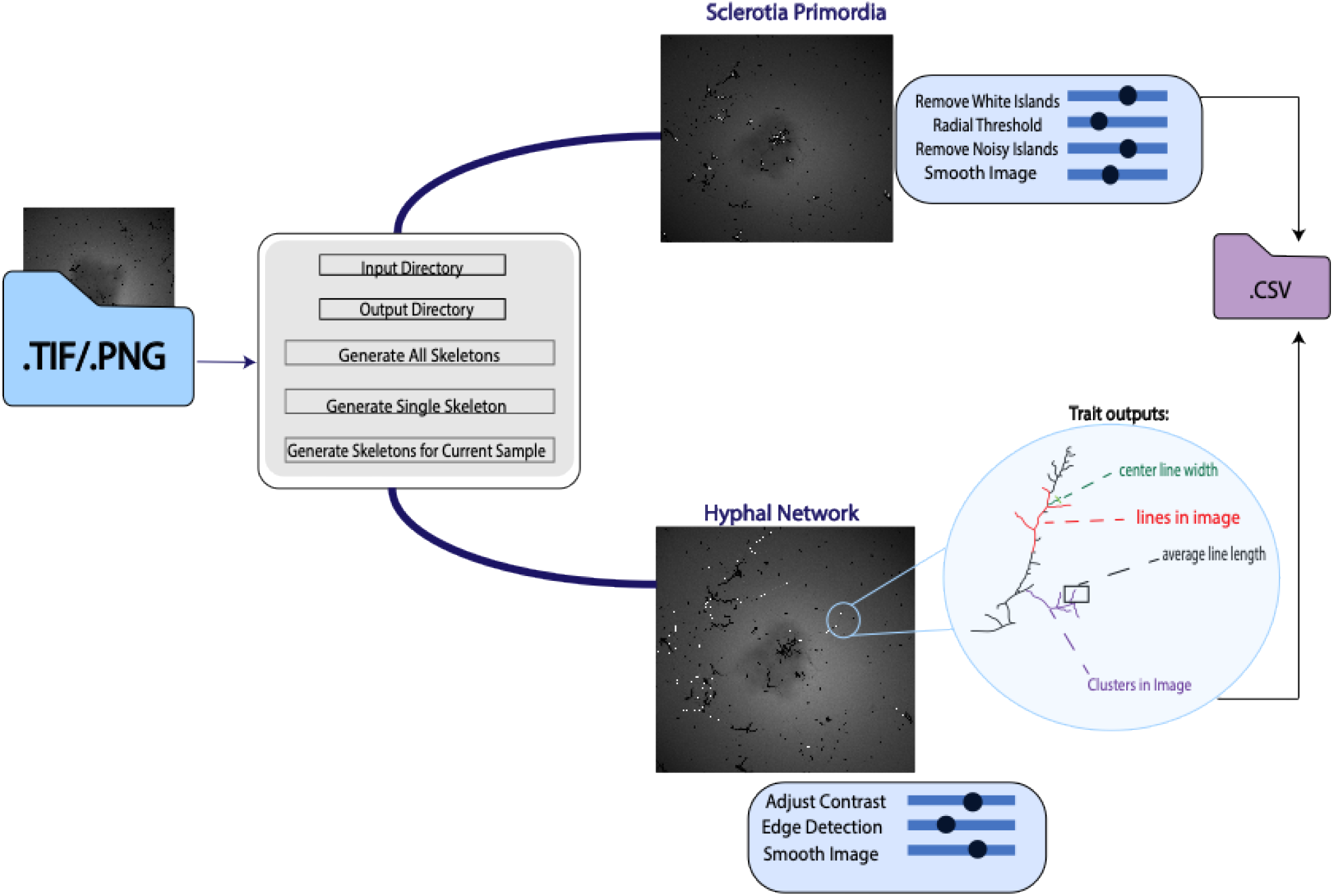
HyPhy Workflow. Users input grayscale images of interest, which are skeletonized and processed to extract network traits such as lines in Image, average line length, and center line width. Parameters for skeletonization can be modified by users, allowing for optimization of different image types. Outputs include skeletonized images and CSV files containing coordinates and trait data for downstream analysis.

### Image Processing

Step-by-step instructions for using HyPhy are available on GitHub and described here. To run HyPhy, users can run the following code in their terminal: python MainApplication.py. HyPhy is made with PyQt, a commonly used library for Python, that establishes communication between parent and child objects. For example, modifying the contrast of the image will send a signal back to the parent object and triggers the corresponding image-processing function.

Skeletons of the images are generated by going through radial threshold, which creates a threshold mask that interpolates from the centerThreshold at the center of the image to edgeThreshold at the edges of the image. This returns an input image thresholded pixel-wise with the threshold mask. The thresholded image removes any remaining white blobs in the image that are fewer than the minWhiteIslandSize pixels threshold set by the user. This step prevents any black blobs from being included in the output.

These blobs create noise, which we define as pixels that are disconnected from our hyphal network or sclerotia primordia. To reduce background noise, pixels that averaged more than four black neighbors were removed, as these are representative of the noise in the image. Neighbors are defined as the eight surrounding positions within a 3×3 pixel window. The number of neighbors per white pixel ranges from zero (pixel is fully embedded within the white region) to eight (pixel is isolated within the black background). The average value is taken across all pixels in the connected component, where a lower value reflects contiguous hyphal networks, and a higher value reflects isolated noise.

The image is then smoothed using gaussian blurring, this ensures that lines of the hyphal network that are connected are represented as connected. The contrast of the image can be modified by using the slide scale, which moves pixel values away or toward 0.5. Adjusting the contrast makes it easier to spot the network lines. To finalize the skeleton for the hyphal network, canny edge detection was applied to identify pixels with strong intensity relative to their neighbors. To reduce noise, a pixel was retained as part of the skeleton if 50% of the pixels within a 2-pixel radius are marked as an edge. This filtering step is necessary because canny edge detection generates a boundary around the skeleton rather than the centerline of the structure. The preprocessed image is then binarized to form a skeleton using Zhang’s skeletonization algorithm (Zhang & Suen, 1984).

After running the application, the skeleton viewer allows users to view the input and vectorized skeletonized image side by side. Users can overlay the skeleton on the original image and change the preprocessing steps if needed and preview them. The preview window is accessed when the user clicks on the preview steps button in the main window. The preview window displays the current skeleton, input image, the step the user is currently modifying, all the parameters associated with the current step, and sliders for the parameters. By clicking the left and right buttons, user can manually step through the skeletonization process for every skeleton. If parameters are adjusted, they can be refreshed by clicking the refresh button and previewing the output skeleton. Users can modify the skeletonization pipeline and create their own skeletonization functions in Python for images of interest. HyPhy extracts two skeletons by default, the first skeleton segments the sclerotia primordia and the second segments the hyphal network.

### Trait Extraction in GUI

From the generated skeletons, a wide variety of traits can be extracted (**Table 2**). The GUI’s built-in traits include fractal dimension, which has been previously demonstrated to be a useful metric by (Aguilar-Trigueros et al., 2022; Vidal-Diez De Ulzurrun et al., 2015).

**Table 2.**
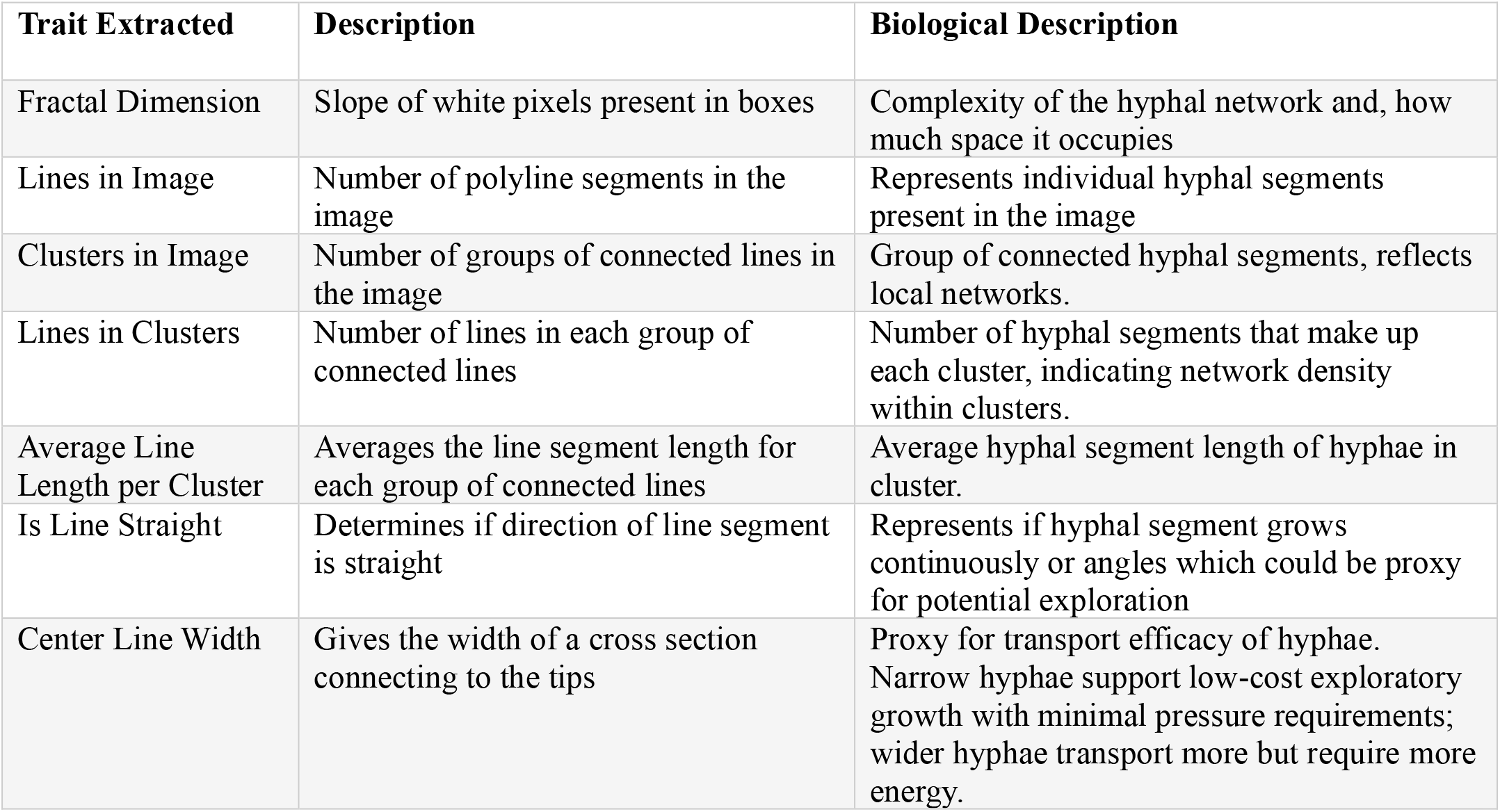
Trait Descriptions.

| Trait Extracted | Description | Biological Description |
| --- | --- | --- |
| Fractal Dimension | Slope of white pixels present in boxes | Complexity of the hyphal network and, how much space it occupies |
| Lines in Image | Number of polyline segments in the image | Represents individual hyphal segments present in the image |
| Clusters in Image | Number of groups of connected lines in the image | Group of connected hyphal segments, reflects local networks. |
| Lines in Clusters | Number of lines in each group of connected lines | Number of hyphal segments that make up each cluster, indicating network density within clusters. |
| Average Line Length per Cluster | Averages the line segment length for each group of connected lines | Average hyphal segment length of hyphae in cluster. |
| Is Line Straight | Determines if direction of line segment is straight | Represents if hyphal segment grows continuously or angles which could be proxy for potential exploration |
| Center Line Width | Gives the width of a cross section connecting to the tips | Proxy for transport efficacy of hyphae. Narrow hyphae support low-cost exploratory growth with minimal pressure requirements; wider hyphae transport more but require more energy. |

In HyPhy **fractal dimension** serves as a description of how complicated the network in the image represents. It is calculated using a box-counting method in which a box is created and walked across an image. This box counts how many times a white pixel is present. The box size increases as it walks through the image. These values are then used to calculate the slope of the number of boxes that contain white pixels. Another useful trait is the **clusters in image**. It represents the number of groups of connected lines in the image, like hyphal segments that have branched and connected, forming interconnected networks rather than isolated filaments. All traits are calculated in Python functions, to which the user can modify and expand. Together, these traits give us better insight into the foraging strategies whether fungi are maximizing growth through outward expansion of simple hyphae threads or fungi that invest in branching and form more robust networks.

Another trait is the number of **lines in the image** (continuous growth between branch points). **Lines in clusters** counts the number of lines in each group of connected lines, which measures the hyphal segments that groups are composed of. **Width** of hyphal segments are calculated by finding the center point of the line, getting the direction of the line and its orthogonal direction, and counting the white pixels in the orthogonal line. Width serves as a proxy for transport efficacy.

To support scalability, traits are exported into sample specific folders, minimizing memory demands when working with large datasets. Metadata for each network is saved in network or sclerotiaPrimordiametadata.csv. If additional skeletonization pipelines are implemented, each will get its own CSV and JSON outputs, named according to pipeline. Coordinates and indices for the hyphal network are also provided and can be used for downstream analysis in R.

### R Analysis: Hyphal Coordinates

The coordinates of the hyphal network were processed using the R package igraph to calculate edges, tips, branches, and isolated points that represent ungerminated spores. Edges are described as links that connect points, represent individual hyphal segments and provide a measure of network connectivity. Tips are endpoints of a hyphal segment and serve as a proxy of hyphal extension. Branches occur when a hyphal segment splits to give rise to another hyphal segment which reflects network expansion. The codes used to run this analysis can be found at https://github.com/melaniemadrigal07/HyPhyAnalysis.git.

### Validation of HyPhy

Images of 1.04.25’s hyphal network were hand traced on an iPad Air 3^rd^ generation (Apple, Cupertino, California, USA) using the Notability app (Ginger Labs, Redwood City, California). Sclerotia primordia were traced using pen size 3 and hyphal networks were traced using pen size 1. These images were then skeletonized in ImageJ by using the color adjust threshold, making them binary, and using the skeletonize function to output skeletonized images. These images were inputted into Hyphy’s comparison function to obtain the maximum and average farthest distance to the nearest point.

The modified gridline method to calculate colonization rate of mycorrhizal fungi was used to obtain the colonization rate of our fungal images (Stahlhut et al., 2021). In this method, ten vertical lines were superimposed on the dataset of images. For each image, a single experienced user scored presence/absence of a hyphal segment (1 if hyphae touched the line; otherwise, 0). Colonization was calculated using the following formula: (*Number of lines with intersections*)/(*Total number of lines*) × 100. To validate these measures against Hyphy, we performed Pearson’s correlation between (i) colonization rate and fractal dimension and (ii) hyphal length density and linesinimage.

HyPhy allowed us to skeletonize 180 images on an external hard drive. It took 102 minutes on an Apple M4 MacBook pro 2024 and used 234 MB. To assess performance at scale, we applied HyPhy on a larger dataset of 3,842 images. This larger dataset took 12 hours and 30 minutes. For comparison, hand tracing the sclerotia primordia and hyphal network was substantially slower: an early timepoint image with minimal branching required 23 minutes and 19 seconds, while a later timepoint with extensive branching took approximately 2.5 hours. HyPhy skeletonized those images in less than a minute. To validate automated results, we used HyPhy’s image overlay feature to compare hand-traced skeletonized images and the automated skeleton. On average, automated and hand-traced skeletons differed by 0.0046 units (SD=0.0009), while the maximum distance to the closest point was 0.0195±0.005.

We validated the accuracy of HyPhy by using the grid-line method on skeletonized images to estimate hyphal length density. Comparison with the linesInImage metric revealed a strong correlation of (r = 0.99; **Fig. 2**). Grid-line method derived colonization rate was strongly correlated with fractal dimension (r = 0.81;**Fig. 3**). These results indicate that HyPhy derived traits successfully capture variation in fungal network growth. HyPhy derived traits also capture more variation than the colonization method as once 10 intersections were made on the vertical lines the space was considered fully colonized (**Fig. 3**).

**Figure 2.**
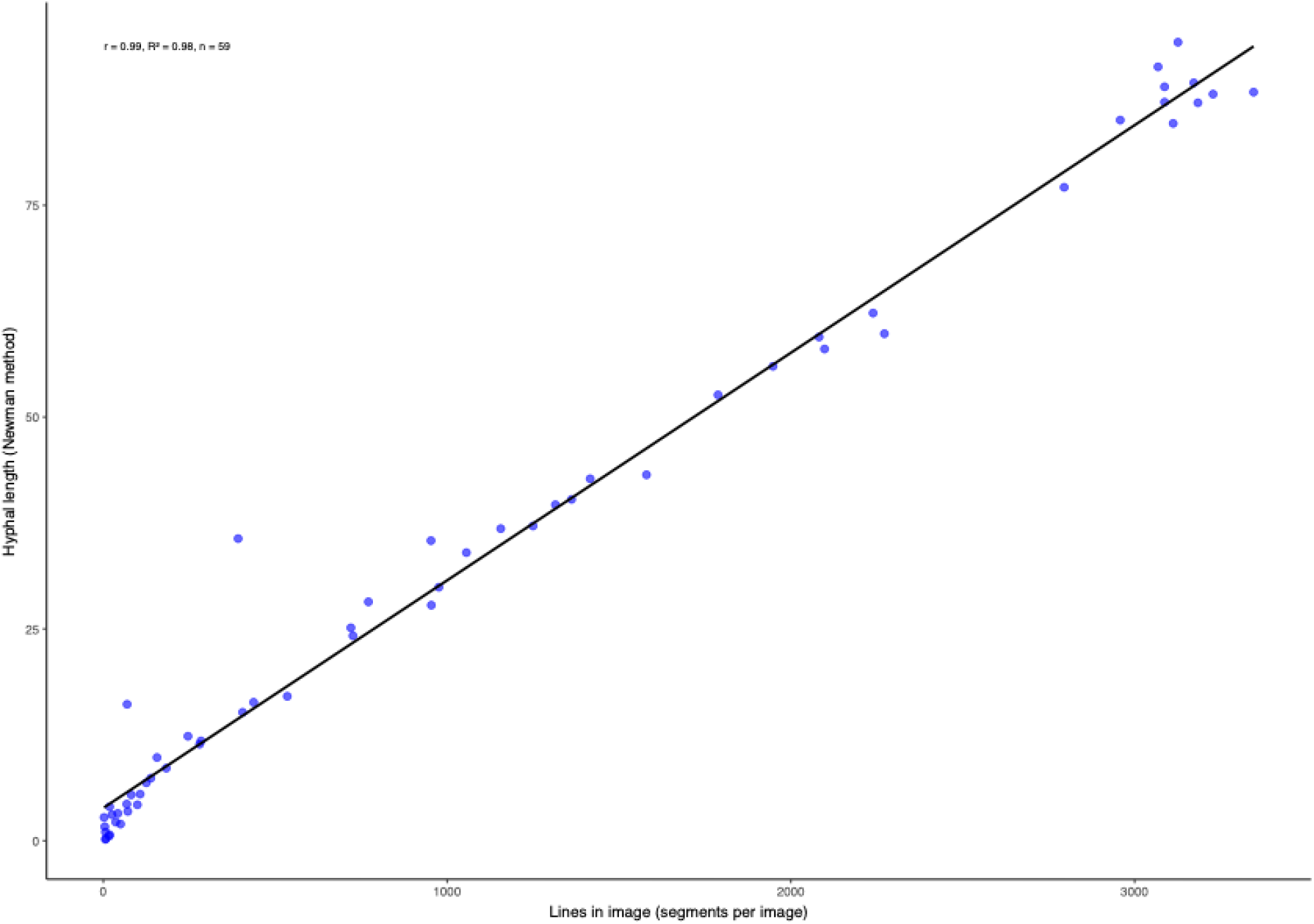
Hyphal Length estimates from the Newman method are strongly correlated with the linesInImage metric (r = 0.99)

**Figure 3.**
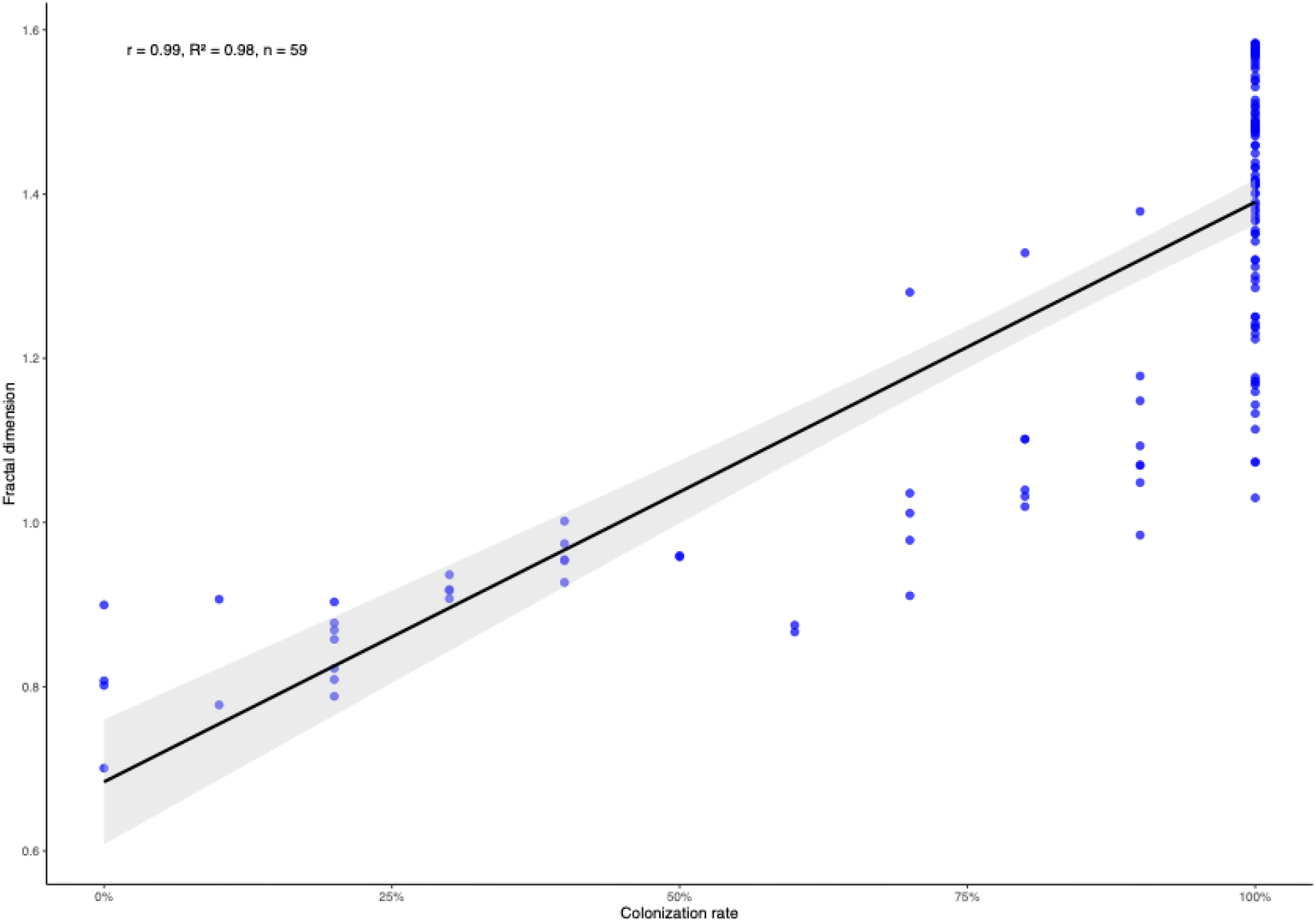
Colonization rate derived from the grid-line method was also strongly correlated with fractal dimension (r = 0.81).

Inputting the coordinates into igraph, we extracted the number of tips, edges, and branches for each strain and we distinguished among isolates with different growth strategies (P = 1.23 ×10^−8^).

KatieTomato, B05.10, and 1.04.25 displayed distincit network pattherns that reflect their differences in virulence. KatieTomato formed the most expansive networks with a maximum of 1,848 active tips, 4,819 edges, and 504 branching events. These traits are consistent with its highly aggressive infection strategy and large lesion formation on hosts. However, its fractal dimension increased at the slowest rate of (0.0232) indicating that despite rapid proliferation, its overall network complexity occurred at a gradual pace. B05.10 represented an intermediate response with a maximum of 1,213 tips, 3,757 edges and 204 branching events. Although B05.10 did not display aggressive proliferation, its fractal dimension increased at the steepest rate 0.0519±0.00326. Despite its lower proliferation its network complexity accumulated at a faster rate, suggesting a growth strategy where a complex network is preferred over maximizing the number of tips and branches. 1.04.25 the slower and less virulent isolate had a maximum of 958 tips, 2,698 edges, and 148 branching events. Its complexity increased at an intermediate rate 0.0398±0.00305, consistent with its phenotype.

## CONCLUSIONS

We provide an accessible user-based Python interface, that extracts meaningful data from hyphal network images that can be used to study variation in fungal morphology. This interface allows users to customize skeletonization parameters to maximize the detection of the hyphal network. HyPhy handles large image datasets and is a powerful method for analyzing hyphal growth. The export of coordinates of the hyphal network allows for the extraction of more traits using network theory such as edges, nodes, and tips that have been previously extracted in other hyphal network extraction programs (Vidal-Diez de Ulzurrun et al. 2015; Aguilar-Trigueros et al., 2022). This interface saves time for users as a complex network image requires approximately two hours, when scaled to datasets of thousands of images, the time savings become substantial. By streamlining these processes our interface makes large-scale analyses feasible, giving researchers more room to focus on the biological interpretation than on a collection. Importantly, we demonstrate that HyPhy can distinguish among isolates of B. cinerea with varying levels of virulence and genetic diversity. Linking quantitative network traits such as tips, branching events, and fractal growth rates to known pathogenetic strategies. This highlights the ability of HyPhy to connect structural traits with biological outcomes, creating a foundation for further modeling approaches.

The use of the extracted traits (Table 2) can be expanded to the utilization of exponential family random graph models (ERGM). ERGM models describe the selection forces that shape the global structure of a network (Hunter et al., 2008). These models could be used to describe structure and pattern within the networks and the selection process that guides hyphal architecture in response to different environmental changes. ERGM models have been used with *E. coli* to identify operons that encode for transcription factors that regulate the second operon (Saul & Filkov, 2007). A similar approach could be used with the hyphal network, to identify if any growth patterns are associated with tested environments. For example, using the graphed skeletons we could fit an ERGM model to identify motifs that could hint at the growth strategy an isolate used by looking at overall connectivity of edges. Models could be compared across treatments to test whether exposure to other compounds shifts hyphal architecture to more exploratory or invasive.

Additionally, the skeletonization function of HyPhy can be modified for diverse experimental designs, it allows users to add or remove skeleton pipelines as needed. For example, if a fluorescent stain was used in the image a custom pipeline could be introduced to isolate stained regions prior to skeletonization to extract structures of interests. HyPhy’s modular design makes this process straightforward, allowing researchers who might not be experienced with image analysis to easily separate target components for downstream structural analysis. By enabling users to customize inputs and tailor the frameworks being modeled, HyPhy provides an accessible platform for adapting network-based approaches to a wide range of biological imaging applications.

## Acknowledgements

J.D., M.M., and J.M. were funded in part by a USDA grant (2022-67012-43019). M.M. was also funded by the Future scholar’s fellowship at Louisiana State University.

## Author Contributions

M.M. and J.A.D. designed study; M.M. and J.C.M. collected the data; A.J.M. and M.M. developed HyPhy and its extensions; M.M, J.C.M., and J.A.D. analyzed data; and M.M. and J.A.D. wrote the manuscript with input from A.J.M. and J.C.M.

## Data Availability Statements

The HyPhy source code, and example images used to demonstrate the pipeline, are available at. https://github.com/AaronMoseley/HyPhy.git. Code for downstream analyses in R, including data preparation and trait extraction, is available at https://github.com/melaniemadrigal07/HyPhyAnalysis.git.

